# Association between premorbid body mass index and amyotrophic lateral sclerosis: causal inference through genetic approaches

**DOI:** 10.1101/526186

**Authors:** Ping Zeng, Xinghao Yu, Haibo Xu

## Abstract

**Background:** Inverse association between premorbid body mass index (BMI) and amyotrophic lateral sclerosis (ALS) has been discovered in observational studies; however, whether this association is causal remains largely unknown.

**Methods:** We employed a two-sample Mendelian randomization approach to evaluate the causal relationship of genetically increased BMI with the risk of ALS. The analyses were implemented using summary statistics obtained for the independent instruments identified from large-scale genome-wide association studies of BMI (up to ∼770,000 individuals) and ALS (up to ∼81,000 individuals). The causal relationship between BMI and ALS was estimated using inverse-variance weighted methods and was further validated through extensive complementary and sensitivity analyses.

**Findings:** Using 1,031 instruments strongly related to BMI, the causal effect of per one standard deviation increase of BMI was estimated to be 1.04 (95% CI 0.97∼1.11, *p*=0.275) in the European population. The null association between BMI and ALS discovered in the European population also held in the East Asian population and was robust against various modeling assumptions and outlier biases. Additionally, the Egger-regression and MR-PRESSO ruled out the possibility of horizontal pleiotropic effects of instruments.

**Interpretation:** Our results do not support the causal role of genetically increased or decreased BMI on the risk of ALS.

## 1. Introduction

Amyotrophic lateral sclerosis (ALS) is a frequent adult-onset fatal neurodegenerative disease clinically characterized by rapidly progressive motor neurons degeneration and death because of respiratory failure [1]. Although great advance has been made for the understanding of ALS in the past decades, the pathogenic mechanism underlying ALS remains largely unknown and only few therapeutic options can be available [2]. It has been shown that both genetic [3, 4] and environmental factors (e.g. cigarette smoking, alcohol consumption, exposure to pesticides, lead, organic toxins or electromagnetic radiation and socioeconomic status) may contribute to the development of ALS [5, 6, 7, 8, 9, 10, 11]. However, no replicable and definitive environmental risk factors are currently well established for ALS. In addition, due to the quickly growing ageing of the population in the upcoming years, it is evaluated that the number of ALS cases across globe will increase by about 70% [12], which is anticipated to result in rather serious socioeconomic and health burden. Therefore, understanding the risk factors of ALS for improving the medical intervention and quality of life for ALS patients is considerably important from both disease treatment and public health perspectives.

Among extensive epidemiological researches of ALS, an interesting and surprising observation is that ALS patients often encounter a loss of weight or a decrease of body mass index (BMI) at the early phase of diagnosis with many possible explanations [1, 13, 14, 15, 16, 17, 18, 19, 20]. Indeed, substantial change of BMI in ALS patients has been identified as an independent prognostic factor and has been linked to disease progression [13, 21, 22, 23, 24, 25, 26, 27]. For example, it is observed that a rapid reduction of BMI in ALS patients at the initial disease stage is a strong indicator of faster disease progression and shorter survival time; in contrast, nutritional intervention for ALS patients to increase BMI can prolong the survival time and leads to a delay in disease progression [28, 29, 30]. The benefit of raising BMI for ALS patients by taking high-energy diet is also confirmed by mouse models [31].

However, whether the long-term exposure to genetically increased (or decreased) BMI prior to the onset of ALS also plays a pathological role in the development of ALS is less understood. Various findings with regard to the relationship between premorbid (i.e. prediagnostic) BMI and ALS have been reported in the literature. In a large-scale observational cohort, it was shown that higher BMI before the onset of ALS was associated with a decreased risk of ALS, resulting in an average of 4.6% [95% confidence interval (CI) 3.0∼6.1%] lower risk of ALS per unit increase in BMI [23]. This inverse association between premorbid BMI and the risk (or mortality) of ALS was also supported by some recent studies [32, 33, 34, 35, 36]; see Tables S1 for more information. However, contradictory results were also reported. For example, it was shown that ALS cases consistently had a greater BMI compared with controls beyond five years before ALS manifestation although they had a smaller BMI than controls within five years prior to onset; and that per unit increase of BMI can result in ∼5.0% (95% CI 0.0∼11.0%) higher risk of ALS (Table S2 and Fig. S1) [13]. This study also implied that BMI may already begin to change about ten years before onset of ALS. A more pronounced increased risk associated with greater BMI before five years onset of ALS was observed in a population-based case-control study performed in Washington State [37, 38]: 50% higher risk of ALS for those with BMI between 24 and 26 kg/m^2^ compared with those with BMI less than 21 kg/m^2^, and 70% higher risk of ALS for those with BMI between larger than 26 kg/m^2^ compared with those with BMI less than 21 kg/m^2^.

The conflicting observations on the relationship between premorbid BMI and ALS may be partly due to uncontrolled/unknown bias or confounding factors that are frequent in observational studies, or partly due to the relatively small sample size for patients because of the rarity of ALS, or partly due to the reverse causality in previous studies as well as a limited retrospective time (or follow-up) before ALS onset [13]. We employ the systematic review and meta-analysis to provide a pooled conclusion about the relationship between premorbid BMI and ALS (Supplementary Text S1). We show that on average a unit increase of premorbid BMI can robustly result in about 3.0% (95% CI 2.1∼4.5%) risk reduction of ALS, supporting the previous finding that a greater premorbid BMI is a protective factor for the development of ALS [23, 32, 33, 34, 35, 36]. Nevertheless, there still exists an essential problem — is the change of BMI before ALS manifestation a causal risk factor or the consequence of ALS?

Because BMI is a modifiable exposure factor and obesity is a growing global health problem [39], a better characterization of the causal effect of BMI on ALS can thus facilitate our understanding of the pathogenesis of ALS and finally leads to better prevention and treatment for ALS patients. Traditionally, randomized controlled trial (RCT) studies are the gold standard for inferring the causal impact of exposure on outcome. However, determining the causal relationship between premorbid BMI and ALS through RCT is challenging and unrealistic, because RCT necessarily requires a very large set of subjects and an extremely long follow-up before clinical manifest of ALS due to its rarity in the population [40] and wide variations in prevalence and incidence across various age groups [41, 42, 43, 44]. Therefore, it is desirable to investigate the causal association between premorbid BMI and ALS through observational studies. Mendelian randomization is such an approach [45, 46], which employs single nucleotide polymorphisms (SNPs) as instruments for the exposure (i.e. premorbid BMI) and assesses its causal effect on the outcome of interest (i.e. ALS) (Fig. S2) [47]. The recent successes of large-scale genome-wide association studies (GWASs) [48, 49, 50, 51, 52] make it feasible to choose strongly associated SNPs to be valid instruments for causal inference in Mendelian randomization [53, 54]. Indeed, in the last few years Mendelian randomization has become a very popular method for causality inference in observational studies [54, 55].

Therefore, our main goal in this study is to examine whether there exists a causal association between the long-term exposure to genetically increased (or decreased) BMI and ALS onset. Consequently, SNPs which influence BMI would also affect the risk of ALS through changes of BMI. To do so, we conducted the largest and most comprehensive two-sample Mendelian randomization analysis to date by using summary statistics obtained from large-scale GWASs with ∼770,000 individuals for BMI and ∼21,000 ALS cases.

## 2. Materials and Methods

### 2.1. GWAS data sources and instrument selection

We selected 1,031 independent index association SNPs (*p*<5.00E-8) to serve as instrumental variables for BMI (Table S3) from the Genetic Investigation of ANthropometric Traits (GIANT) consortium, which is the largest BMI GWAS (up to 773,253 individuals) for the European population to date (Supplementary Text S2) [50]. For all the instruments we obtained their association summary statistics in terms of the effect allele, marginal effect size estimate and standard error. To estimate the causal effect of BMI on ALS, we extracted corresponding association summary statistics of these index SNPs for ALS from an ALS GWAS that was also carried out in the European population up on 80,610 individuals (20,806 cases and 59,804 controls) (Table S3) (Supplementary Text S2) [4].

Besides the set of 1,031 instruments obtained from [50], as a part of complementary and sensitivity analyses, we also attempted to validate whether the relationship between BMI and ALS derived from the European population also holds in the East Asian population. To do so, we performed an additional Mendelian randomization study using another set of 75 instruments obtained from an East Asian BMI GWAS up to 158,284 individuals (Table S4 and Supplementary Text S2) [52]. The corresponding summary statistics of ALS for these instruments were extracted from an East Asian ALS GWAS up to 4,084 individuals (1,234 cases and 2,850 controls) (Supplementary Text S2) [56]. The two sets of index SNPs of BMI from the two populations share only one common instrument (i.e. rs7903146). The GWAS data sets used in the present study are summarized in Table 1.

**Table 1.**
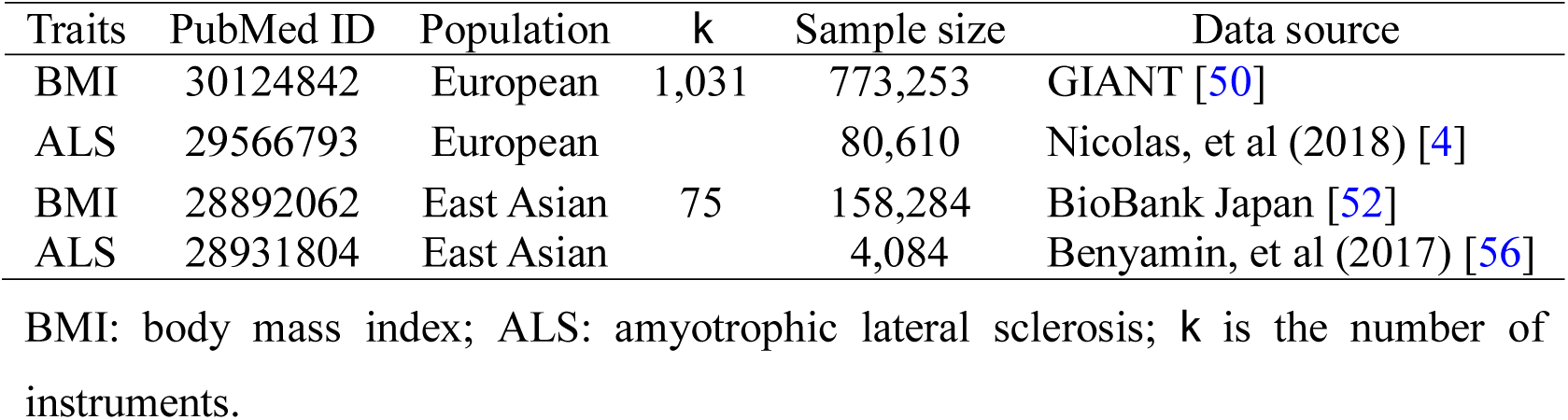
GWAS data sets used in the Mendelian randomization analysis in the main text. Traits

| Traits | PubMed ID | Population | $k$ | Sample size | Data source |
| --- | --- | --- | --- | --- | --- |
| BMI | 30124842 | European | 1,031 | 773,253 | GIANT [50] |
| ALS | 29566793 | European |  | 80,610 | Nicolas, et al (2018) [4] |
| BMI | 28892062 | East Asian | 75 | 158,284 | BioBank Japan [52] |
| ALS | 28931804 | East Asian |  | 4,084 | Benyamin, et al (2017) [56] |

BMI: body mass index; ALS: amyotrophic lateral sclerosis; $k$ is the number of instruments.

### 2.2. Estimation of causal effect with inverse-variance weighted methods

To examine whether the instruments are strong, for each index SNP that was used as instrument in turn, we calculated the proportion of phenotypic variance of BMI explained (PVE) by the instrument using summary statistics [57] and generated the *F* statistic (Table S3) [58, 59]. We then performed the two-sample Mendelian randomization analysis [60, 61] to estimate the causal effect of BMI on ALS in terms of OR per standard deviation (SD) change in BMI with inverse-variance weighted (IVW) methods [59, 62]. Before formal analysis, to ensure the validity of Mendelian randomization analysis, we examined the pleiotropic associations of instruments by removing those that may be associated with ALS with a marginal p value below 0.05 after Bonferroni correction. While in our analysis no instruments were excluded from any set of instruments by this strategy.

### 2.3. Complementary and sensitivity analyses

To ensure results robustness and guard against model assumptions in the Mendelian randomization analysis (Fig. S2), we carried out a series of complementary and sensitivity analyses: (**i**) leave-one-out (LOO) cross-validation analysis [63] and Mendelian Randomization Pleiotropy RESidual Sum and Outlier (MR-PRESSO) analysis [64] to validate whether there are instrumental outliers that can substantially influence the causal effect estimate; (**ii**) weighted median-based method that is robust when some instruments are invalid [65]; (**iii**) MR-Egger regression to examine the assumption of directional pleiotropic effects [61, 66]; (**iv**) IVW causal analysis after removing instruments that may be correlated to other 38 complex metabolic, anthropometric and socioeconomic traits from large-scale GWASs (Supplementary Text S2); (**v**) reverse causal inference on BMI using ALS instruments; (**vi**) IVW method for the causal effect estimation of BMI on ALS in the East Asian population.

## 3. Results

### 3.1. Causal effect of BMI on ALS

By using the PLINK procedure (Supplementary Text S2), we selected 1,031 independent SNPs to be valid instruments for BMI in the European population from the GIANT study [50] (Table S3). These instruments together explain a total of 8.28% of phenotypic variance for BMI. The *F* statistics for all these SNPs are above 10 (ranging from 28.8 to 1426.2 with an average of 58.4), implying that the weak instrument is less likely to bias our analysis. In addition, as there is significantly statistical evidence for the heterogeneity of causal effect across instruments (*p*=7.43E-4); therefore, only the results estimated using the random-effects IVW method are displayed in the following paragraphs.

With the random-effects IVW method we find that the OR of ALS per unit [one standard deviation (SD)] increase of BMI is estimated to be 1.04 (95% CI 0.97∼1.11, *p*=0.275) using the set of 1,031 instruments, suggesting that the genetically changed BMI is not necessarily causally associated with an increased or decreased risk of ALS. We further examine whether the lack of detectable non-zero causal effect of BMI on ALS is due to a lack of statistical power. To do so, we calculated the statistical power to detect an OR ratio of 1.10 (or 0.90) in the risk of ALS per unit change of BMI following the analytic-form approach given in [67] (https://cnsgenomics.shinyapps.io/mRnd/). It is shown that the estimated statistical power is 94%, indicating that we would have reasonably high power to detect such a causal effect of BMI on ALS if BMI is indeed causally related to the risk of ALS.

### 3.2. Sensitivity analyses to validate the estimated causal effect of BMI on ALS

We now validate the causal effect of BMI on ALS estimated above through various sensitivity analyses. First, we examine whether there exist potential instrument outliers and whether these outliers have a substantial influence on the estimate of causal effect. To do so, we created a scatter plot by drawing the effect sizes of BMI with regard to their effect sizes of ALS for all the 1,031 instruments. Among all the instruments, one index SNP (i.e. rs2229616) has the largest effect size of 0.106 on BMI and can be reasonably assumed to be a potential outlier (Fig. 1). However, this outlier does not impact the estimated causal effect in our Mendelian randomization analysis. Specially, after removing rs2229616, the OR of ALS per one SD increase of BMI is 1.04 (95% CI 0.97∼1.11, *p*=0.317), almost the same as that obtained using all the instruments. To further examine whether a single instrument may strongly influence the causal effect of BMI on ALS, we performed a leave-one-out (LOO) Mendelian randomization analysis. Again, the LOO analysis results show that no single instrument can influence the causal effect estimate substantially (Fig. 2). Additionally, we also directly tested whether any instrument is an outlier using MR-PRESSO [64], which shows that no significant instrument outliers exist for the Mendelian randomization analysis at the significance level of 0.05.

**Fig. 1.**
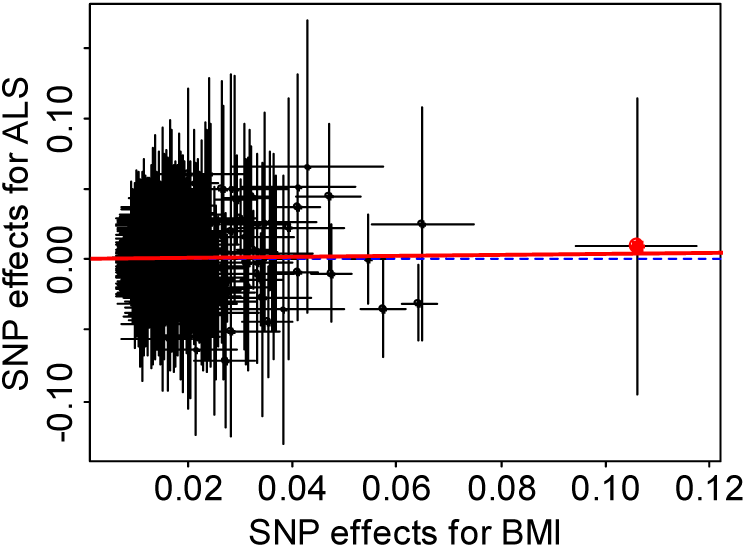
Relationship between the SNP effect size estimates of BMI (x-axis) and the corresponding effect size estimates of ALS (y-axis) in the European population using 1,031 instruments generated from [50]. In the plot, the 95% CIs for the effect sizes of instruments on BMI are shown as horizontal lines, while the 95% CIs for the effect sizes of instruments on ALS are shown as vertical lines. The horizontal dotted line represents zero effects. The line in red represents the estimated causal effect of BMI on ALS obtained using the random-effects IVW method. The red dot in the rightmost side is identified as an outlier (i.e. rs2229616).

**Fig. 2.**
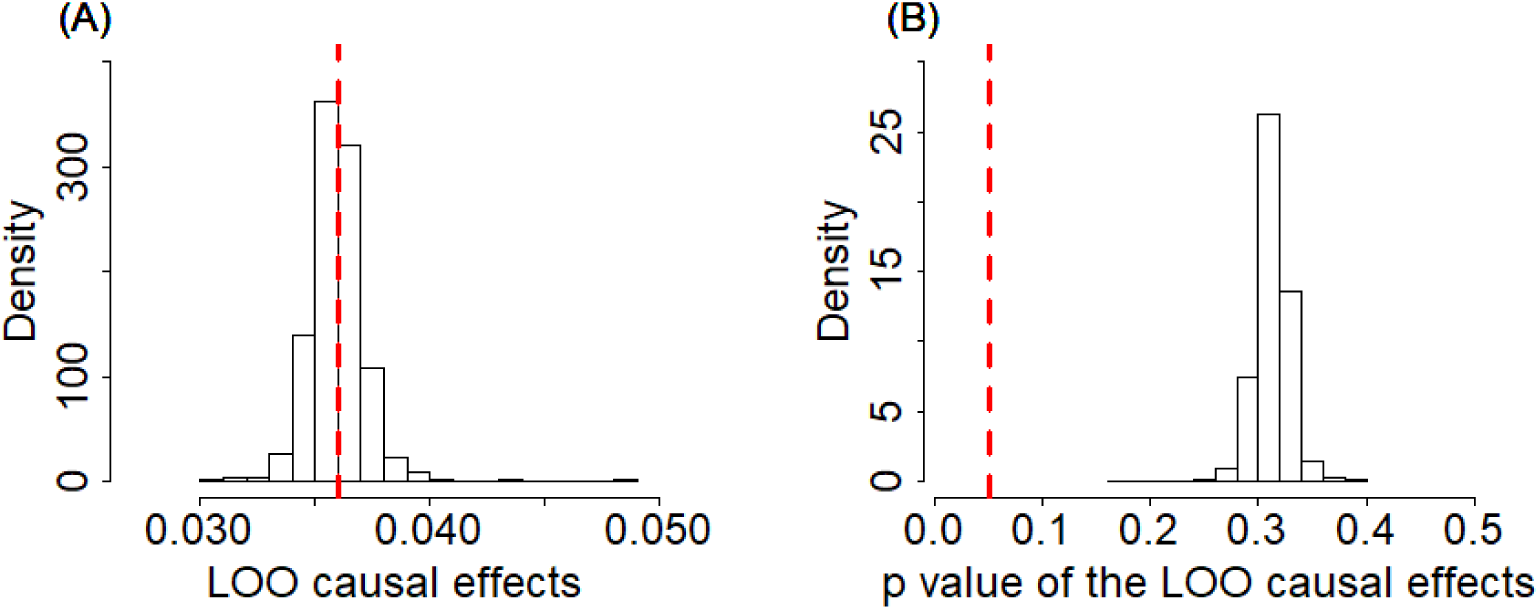
Leave-one-out (LOO) results of BMI on ALS based on the set of 1,091 instruments in the European population. (A) Estimated LOO causal effects; (B) The p values of the LOO causal effects. In the left panel, the red reference line is the point estimate of causal effect for BMI using all the instruments; in the right panel, the red reference line represents the significance level of 0.05.

To study whether some are invalid among the set of 1,031 instruments and may bias the results, we conducted a Mendelian randomization analysis using the weighted median method [65]. The weighted median method yields the consistent estimate as before. In particular, the OR of ALS per one SD increase of BMI is calculated to be 1.01 (95% CI 0.91∼1.13, *p*=0.806), suggesting that invalid instruments unlikely bias our results.

To investigate whether these instruments show potentially horizontal pleiotropy, we performed a Mendelian randomization analysis using the MR-Egger regression [61, 66]. The results from the MR-Egger regression analysis are again largely consistent with our main results. For example, using all the 1,031 instruments the MR-Egger estimates the OR per one SD increase of BMI on ALS to be 0.97 (95% CI 0.79∼1.19, *p*=0.740). The MR-Egger regression intercept is 0.001 (95% CI -0.002∼0.004, *p*=0.473). Furthermore, funnel plots also display a symmetric pattern around the causal effect point estimate (Fig. 3). The funnel plots and MR-Egger regression results together offer no evidence for horizontal pleiotropy.

**Fig. 3.**
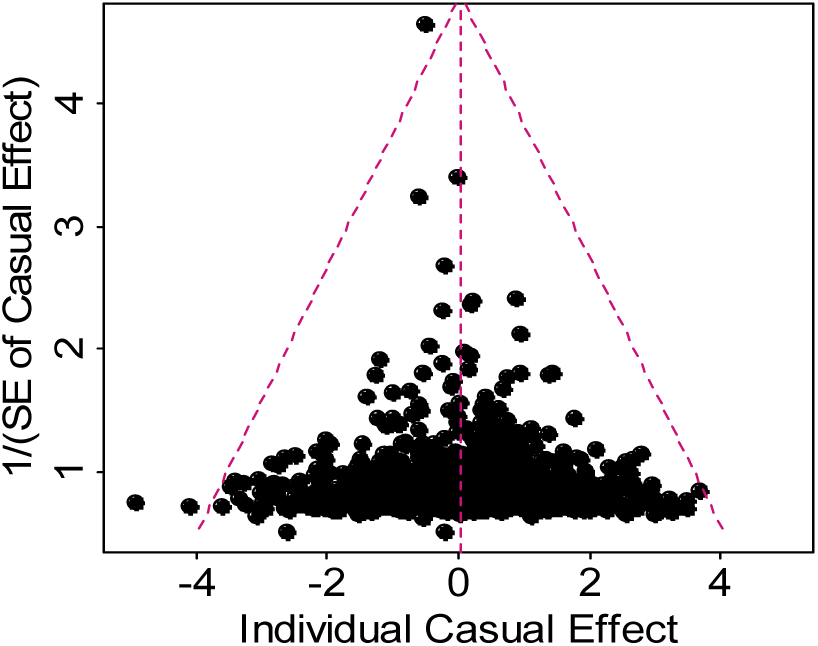
Funnel plot for single causal effect estimate of BMI on ALS obtained using all 1,091 instruments in the European population.

We further performed causal estimation for BMI after removing instruments that may be potentially associated with other 38 complex traits (Supplementary Text S2) with various threshold values. Again, the resulting estimates of causal effect for BMI on ALS are not statistically significant regardless of the thresholds used, consistent with the results obtained with all the instruments (Table S5). The Mendelian randomization analysis using ALS instruments also removes the likelihood of reverse causality (the causal effect size is -0.011 with 95% CI -0.034∼0.011 and *p*=0.317).

Finally, we estimated the causal effect of BMI on ALS in the East Asian population. To do so, we first obtained a set of 75 BMI-associated SNPs to serve as instruments (Tables S4). The estimated PVE by these instruments is 2.59%, and all have an *F* statistic above 10 (range from 22.6 to 410.1 with an average of 56.8) and are thus deemed as strong instruments [58, 59]. With these identified instruments, again we find that genetically higher BMI is not causally associated with an increased or decreased risk of ALS at the significance level of 0.05. Specifically, the OR of ALS per one SD increase of BMI is estimated to be 0.90 (95% CI 0.59∼1.39, *p*=0.647).

## Discussion

The main objective of our study was to dissect whether there has a causal association between premorbid BMI and ALS and to investigate whether genetic predisposition to BMI plays an etiological role in ALS. To achieve this, in the present paper we have implemented a comprehensive Mendelian randomization analysis using summary statistics from GWASs. To our knowledge, this is the first Mendelian randomization study to explore the relationship between BMI prior to disease onset and ALS by leveraging genetic information from large scale GWASs. Additionally, as little has been known about the casual factors for the development of ALS to data[2]; therefore, our study contributes considerably to research on the role of premorbid BMI with regard to the risk of ALS and has the potential implication in public health.

Although previous epidemiological studies showed evidence that premorbid BMI may have a neuroprotective role on ALS (Supplementary Text S1) [32, 33, 34, 35, 36, 37, 38], our Mendelian randomization analysis does not support the existence of causal association between premorbid BMI and the risk of ALS. Besides finding the null causal relationship between BMI and ALS in the European population, we also validated that the causal association does not hold in the East Asian population and that the failure of identifying non-zero causal effect of BMI on ALS is not possibly due to the lack of statistical power in the European population.

Unlike the results of observational studies which can be easily biased by measurement errors and confounding factors (e.g. cigarette smoking, alcohol drinking or daily diet intakes; and it indeed has been shown that BMI is no longer associated with ALS after controlling for socioeconomic status, prior chronic obstructive pulmonary disease, marital status, diabetes and residence at ALS diagnosis [68]), the results of Mendelian randomization analysis are often not susceptible to the measurement error bias because SNPs can be sequenced accurately [69], and are also less susceptible to reverse causation and confounders compared to other study designs because Mendelian randomization uses the principle that the random meiotic assortment of genotypes is independent of confounders and disease process of ALS. Additionally, one of the strengths of this study is that the GWAS data sets with large sample sizes (up to ∼770,000 for BMI [50] and ∼81,000 for ALS [4]) that were employed in our analyses ensures sufficient statistical power and thus the Mendelian randomization results are reasonably believable.

It needs to emphasize that in the main Mendelian randomization analysis we employed multiple uncorrelated strongly relevant instrument variables (a total of 1,031) for causality inference of BMI on ALS. The benefit of applying multiple instruments in Mendelian randomization analysis is that the possibility of weak instruments bias is less likely and the high statistical power is guaranteed. However, it also has a higher likelihood to incorporate pleiotropic instruments, which violates the assumptions of Mendelian randomization analysis (Fig. S2) [47, 53, 54]. Therefore, to minimize the possibility of pleiotropy, we have tried to remove pleiotropic instruments. In addition, we also carried out sensitivity analyses by excluding instruments that may be associated with other 31 complex phenotypes which may link to ALS in a metabolic, anthropometric or socioeconomic way and possibly mediate the effect of BMI on ALS. Our Mendelian randomization analysis showed that the results are robust against pleiotropy and mediation effects as well as various model assumptions.

The mechanisms underlying the causal associations between BMI and ALS may be considerably complex. Although no statistically significant evidence that BMI influences ALS was found in the direct biological pathway in our study, we cannot exclude the probability that BMI can have an impact on ALS via other indirect pathway. For example, it has been extensively shown that BMI is linked to many cardiovascular risk factors (e.g. hypertension and dyslipidemia) and metabolic phenotypes (e.g. type II diabetes, glucose and insulin levels), which in turn have been found to be protective in the development of ALS [70, 71, 72, 73, 74]. The underlying mechanism may be that higher BMI is typically associated with higher blood lipid or glucose levels [39], which could resist against the increased energy consumption and hypermetabolism of ALS patients and thus reduce the hypermetabolic damage on the motor neuron system [17], potentially delaying ALS onset and increasing the survival time [20, 28, 29, 30, 75]. As another explanation, higher BMI was also reported to be associated with higher concentrations of progranulin [76] which could mediate the fat-induced insulin resistance and revert mutant TDP-43 (TAR DNA-binding protein 43) induced axonopathy when overexpression [77], while TDP-43 is widely known to link to the increasing risk of ALS [2]. Therefore, we presume that the observed associations between premorbid BMI and ALS risk in those previous studies could be attributed to the consequence of some unknown biological pleiotropy.

Some limitations of this study should be considered. First, similar to other Mendelian randomization studies, we acknowledge that the validity of our Mendelian randomization relies on some crucial modeling assumptions (Fig. S2) [47, 53, 54], some of which cannot be possible to be fully tested for in the framework of summary-data based Mendelian randomization. Thus, we emphasize that the results obtained in the present study should be interpreted with caution, although we have been extremely careful in selecting instruments to satisfy various model assumptions and have conducted extensive sensitivity analyses to guard against model assumption misspecifications. Second, also like other Mendelian randomization studies, we assumed a linear relationship between BMI and ALS in the Mendelian randomization model; while linearity may be not reasonable in the practice. Thus, we cannot fully rule out the possibility of nonlinear link between BMI and ALS. Third, due to the use of GWAS summary statistics rather than the individual-level data, we cannot test for the causal effect between BMI and ALS stratified by gender or age groups [13, 33].

In conclusion, based the Mendelian randomization results obtained from large-scale GWAS summary statistics, the present study is not supportive of the causal role of genetically increased or decreased BMI on the risk of ALS.

## Acknowledgements

We thank all the GWAS consortium studies for making the summary data publicly available and are grateful of all the investigators and participants contributed to those studies.

## Author contributions

PZ and HX conceived the idea for the study. PZ and XY obtained the data, and performed the data analyses. PZ interpreted the results of the data analyses. All the authors wrote the manuscript.

## Supplementary data

Supplementary File S1 and Supplementary File S2

